# Interaction of N-3-oxododecanoyl homoserine lactone with transcriptional regulator LasR of *Pseudomonas aeruginosa*: Insights from molecular docking and dynamics simulations

**DOI:** 10.1101/121681

**Authors:** Hovakim Grabski, Lernik Hunanyan, Susanna Tiratsuyan, Hrachik Vardapetyan

## Abstract

**Background:** In 2017 World Health Organization announced the list of the most dangerous superbugs and among them is *Pseudomonas aeruginosa,* which is an antibiotic resistant opportunistic human pathogen as well as one of the ‘SKAPE’ pathogens. The central problem is that it affects patients suffering from AIDS, cystic fibrosis, cancer, burn victims etc. *P. aeruginosa* creates and inhabits surface-associated biofilms. Biofilms increase resistance to antibiotics and host immune responses, because of those current treatments are not effective. It is imperative to find new antibacterial treatment strategies against *P. aeruginosa,* but detailed molecular properties of the LasR protein are not clearly known to date. In the present study, we tried to analyse the molecular properties of the LasR protein as well as the mode of its interactions with autoinducer (AI) the N-3-oxododecanoyl homoserine lactone (3-0-C12-HSL).

**Results:** We performed docking and molecular dynamics (MD) simulations of the LasR protein of *P. aeruginosa* with the 3-0-C12-HSL ligand. We assessed the conformational changes of the interaction and analysed the molecular details of the binding of the 3-0-C12-HSL with LasR. A new interaction site of the 3-0-C12-HSL with LasR protein was found, which involves interaction with conservative residues from ligand binding domain (LBD), beta turns in the short linker region (SLR) and DNA binding domain (DBD). It will be referenced as the LBD-SLR-DBD bridge interaction or “the bridge”. We have also performed LasR monomer protein docking and found a new form of dimerization.

**Conclusions:** This study may offer new insights for future experimental studies to detect the interaction of the autoinducer with “the bridge” of LasR protein and a new interaction site for drug design.

## Background

*Pseudomonas aeruginosa (P. aeruginosa*) is a Gram-negative, monoflagellated, obligatory aerobic bacteria [1, 2], This bacterium is one of the ‘SKAPE’ pathogens [3], It can be found in many diverse environments such as soil, plants, hospitals, etc. [1, 4], *P. aeruginosa* is an opportunistic human pathogen, because it rarely infects healthy persons. The central problem is that it affects patients suffering from AIDS, cystic fibrosis, cancer and burn victims [2, 5], Most of the deaths caused by cystic fibrosis are due to this pathogen [1, 6], The pathogenicity of *P. aeruginosa* occurs due to the synthesis of virulence factors such as proteases, hemolysins, exotoxin A, production of antibiotic pyocyanin, Hydrogen Cyanide (HCN), secretion systems of Types 1 (T1SS), 2 (T2SS), 3 (T3SS), 4[7], 5 (T5SS), and 6 (T6SS) [8], ramnolipids and biofilm formation by this organism [7, 9].

Biofilm formation is characteristic to nearly all bacteria where cell-to-cell communication occurs as the population density increases in the human body during pathology. This system is called quorum sensing (QS) that uses hormone-like molecules called autoinducers (AI) that are accumulated in the extracellular matrix. When a threshold is reached, the AIs bind to its cognate receptor and then a response regulator modulates gene expression of QS genes virulence, which includes adaptation, colonization, antibiotic resistance, plasmid conjugation, etc. Among *P. aeruginosa* the AI type 1 system is represented by TuxI/TuxR-typed proteins. AI-1 diffuses freely through the bacterial membrane and binds to the transcriptional activator LuxR. The latter has two LuxI/LuxR-type systems, the first LasI that produces 3-0-C12-HSL and the second, RhlI that synthesizes C4-HSL; both AHLs regulate virulence and biofilm formation. Thus, in *P. aeruginosa* the operon called hcnABC is responsible for HCN biosynthesis through enzyme HCN synthase [10].

Exposure to HCN can lead to neuronal necrosis through the inhibition of cytochrome c oxidase, the terminal component of the aerobic respiratory chain [11, 12], P. aeruginosa has four QS systems [13]. Three transcriptional regulators that perform in a cluster [10] control the transcription of hcnABC genes through LasR, ANR and RhlR [14], The IQS system has been characterized very recently and the genes involved in IQS synthesis are non-ribosomal peptide synthase genes of the ambBCDE operon. The transcriptional regulator of IqsR is the IQS receptor [15].

Nonetheless it was proposed that LasR was the crucial activator of hcnABC genes through mutagenesis experiments [16]. The transcriptional activator protein LasR regulates the target gene expression by recognizing a conserved DNA sequence termed as lux box [16, 17]. LasR has two domains, 1) ligand binding domain at N-terminus (LBD) and 2) DNA binding domain at C-terminus (DBD) [18]. In DBD LasR has a DNA binding Helix-Tum-Helix (HTH) motif [19]. Binding of AI 3-0-C12HSL stabilizes LasR and promotes its dimerization, thereby contributing to the resulting LasR-AI homodimer complex to contact the promoter of the target DNA and activate gene transcription [18].

During colonization and invasion, both the pathogen and the host are exposed to molecules released by each other like bacterial AIs or hosts stress hormones and cytokines. The mechanisms and receptors that might be involved in cross-talk between microbial pathogens and their hosts are not well described to date. LuxR homologues studies have demonstrated that they are homodimers, and consist of two domains. These two functional domains are joined by a short linker region [20, 21].

There remains a need for understanding of the LasR monomer because till date there is no molecular detail information about LasR monomer behaviour. Hence, we analysed the molecular details of the interactions of the 3-0-C12-HSL with LasR protein. So far this is the first report that shows that the 3-0-C12-HSL can interact as well with conservative amino acid residues from ligand binding domain (LBD), beta turns in the short linker region (SLR) and DNA binding domain of LasR. To make easier to reference we refer it as LBD-SLR-DBD bridge or “the bridge”. Could this mean a more nuanced modulation of the transcriptional regulator LasR. This study may be utilized for the development of new therapeutics against *P. aeruginosa* by targeting both the LBD as well as the LBD-SLR-DBD bridge of LasR and inhibit activation of virulence genes.

## Methods

### Analysis of LasR protein sequence

The methodology was based on a previous study [1]. The Amino acid sequence of the LasR protein of *P. aeruginosa* was obtained from UniprotKB (Uniprot id: P25084). The crystal structure of the LBD of LasR protein (amino acid residues 7 to 169) from *P. aeruginosa* was acquired from the Protein Data Bank [22] (PDB id: 3IX3). However, the entire structure of LasR protein was needed in order to have a better understanding of LasR protein properties.

For this reason, the amino acid sequence was used to search in the PDB databank using BLAST [23] to identify suitable templates for homology modelling. The crystal structure of QscR, bound to 3-0-C12-HSL was found to be the best fit (PDB id: 3SZT) [20], The full list of similar structures obtained from BLAST search are shown in Table 1.

**Table 1.**
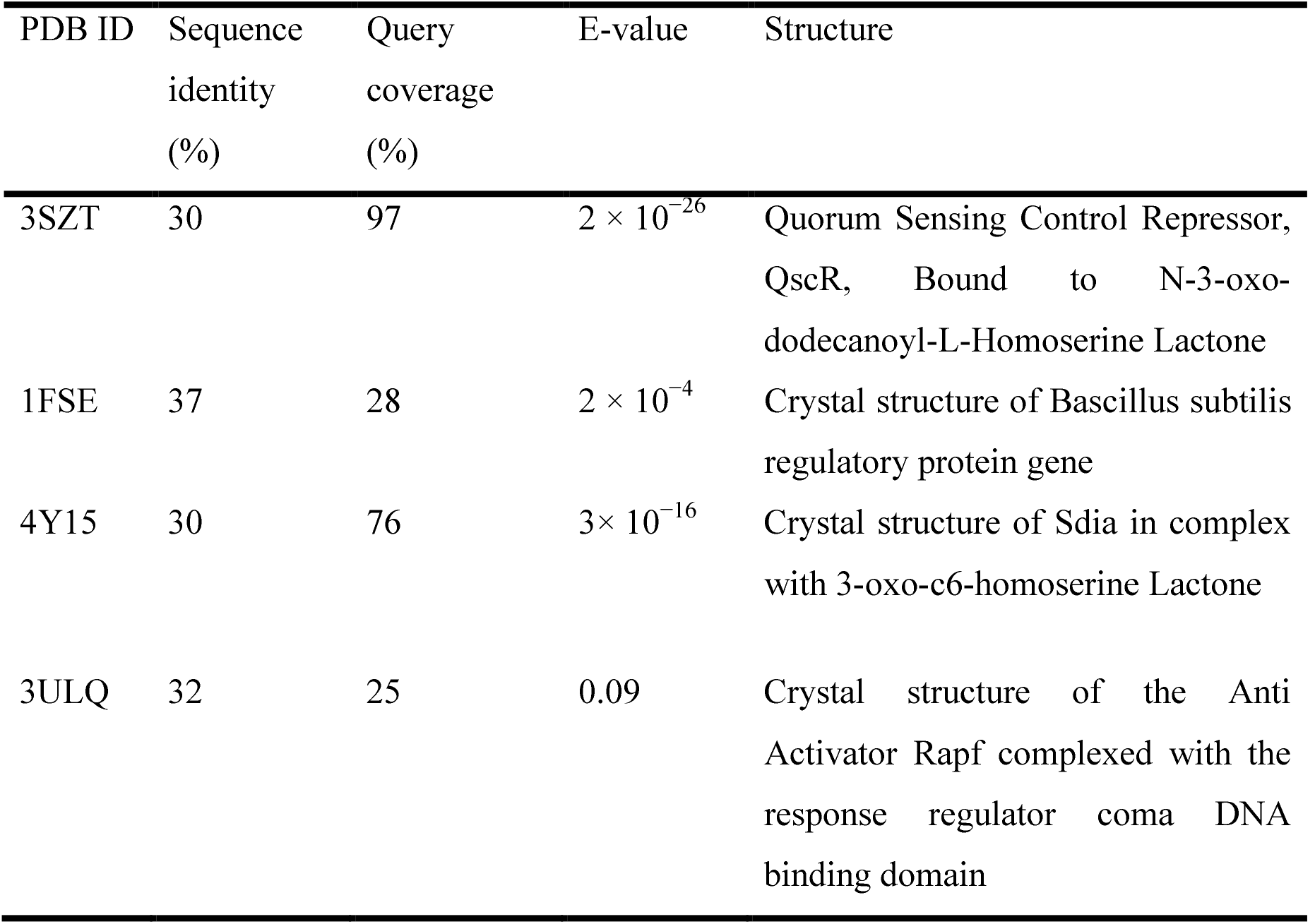
List of similar structures to LasR obtained from BLAST search.

| PDB ID | Sequence<br>identity<br>(%) | Query<br>coverage<br>(%) | E-value | Structure |
| --- | --- | --- | --- | --- |
| 3SZT | 30 | 97 | $2 \times 10^{-26}$ | Quorum Sensing Control Repressor, QscR, Bound to N-3-oxo-dodecanoyl-L-Homoserine Lactone |
| 1FSE | 37 | 28 | $2 \times 10^{-4}$ | Crystal structure of Bascillus subtilis regulatory protein gene |
| 4Y15 | 30 | 76 | $3 \times 10^{-16}$ | Crystal structure of Sdia in complex with 3-oxo-c6-homoserine Lactone |
| 3ULQ | 32 | 25 | 0.09 | Crystal structure of the Anti Activator Rapf complexed with the response regulator coma DNA binding domain |

### Reconstruction of LasR monomer

HHPred web server [24] server was used for the homology modelling of the full LasR protein. The templates used for the homology modelling of LasR are 3SZT, 3IX3, 1FSE and 3ULQ. The aforementioned templates were used to reconstruct the final model of LasR. The final reconstructed model of LasR protein was verified using PROCHECK [25] (Additional file 1: Figures SI, S2), Verify3D [26] and Ramachandran Plots. More than 96.65% of the amino acid residues in the final reconstructed model had a 3D-2D score > 0.2 as indicated by Verify3D as shown in Additional file 1: Figure S3 for a good computational model [26], Ramachandran Plot (Additional file 1: Figure SI) revealed that no amino acid residues were found in the disallowed regions. The PROSA value of this model was -6.69 which suggests that the quality of the homology model is good [27], although the obtained model is slightly different [1], it is still viable.

### Constructing 3-0-C12-HSL model

Figure 1 show the 3D model of 3-0-C 12-FIST, which was acquired from PubChem [28] to study the interactions of the ligand with LasR.

**Fig. 1.**
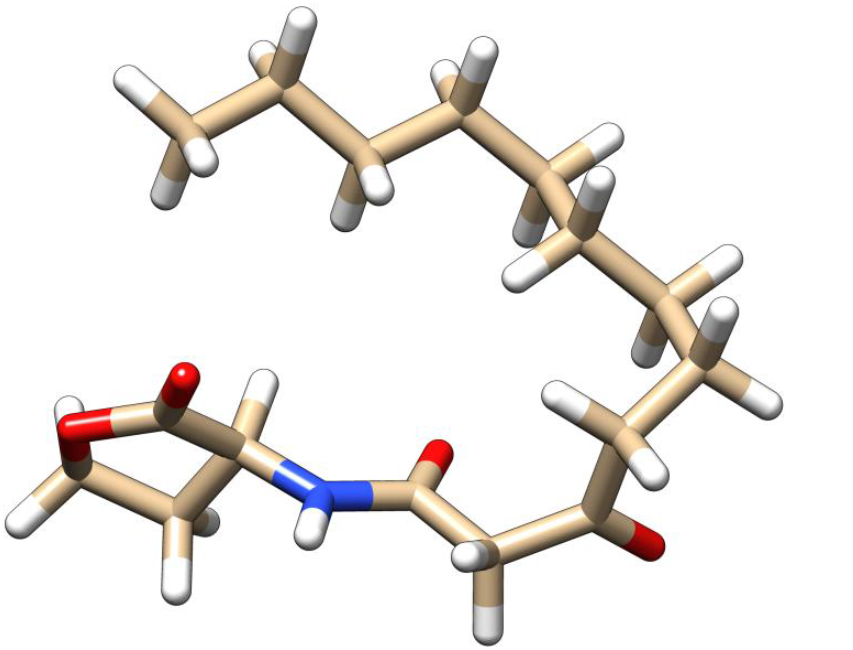
The stick representation of the native ligand of LasR (30-C12-HST).

The ligand parameters for molecular dynamics simulations were calculated for the General Amber Force Field [29] using the acpype tool [30] with AM1-BCC partial charges [31],

### Molecular Dynamics Simulations of LasR protein systems

MD simulations of all systems were conducted with the GROMACS suite, version 5.1.2 [32], utilizing the Amber ff99SB-ILDN force field [33], In all cases, Short-range non-bonded interactions were cut off at 1.4 nm. Particle Mesh Ewald (PME) [34, 35] was used for the calculation of long-range electrostatics.

#### Set 1. LasR monomer simulation

In order to generate the starting structure of LasR monomer before docking, it was placed in a dodecahedron box of TIP3P water [36], to which 100 mM NaCI was added, including neutralizing counter-ions. Following two steepest descents minimization, LasR monomer was equilibrated in two stages. The first stage involved simulating for 200 ps under a constant volume (NVT) ensemble. The second stage involved simulating for 200 ps under a constant-pressure (NPT) for maintaining pressure isotropically at 1.0 bar. Production MD simulation was conducted for 100 ns in the absence of any restraints. The temperature was sustained at 300 K using V-rescale [37] algorithm. For isotropic regulation of the pressure, the Parrinello-Rahman barostat [38] was used.

#### Set 2. Molecular dynamics simulations using docked structures

It has been shown that docking has its limitations [39]. For this reason, after finishing molecular docking simulations we chose the structures of TasR bound to 3-0-C12-HST to perform molecular dynamics (MD) simulations. A time step of 2 fs was used during heating and the production run and coordinates were recorded every 1 ps. Three simulations of 300 ns were performed. Further details about the simulation protocol can be found above.

### LasR-3-0-C12-HSL ligand blind docking experiments

To build the LasR-ligand complex, the ligand 3-0-C12-HST was docked with TasR monomer using Autodock Vina [40]. However, AutoDock Vina currently uses several simplifications that affect the obtained results. The most notable simplification as the creators’ note [40] is the use of a rigid receptor. Vina provides a parameter called ‘Exhaustiveness’ to change the amount of computational effort used during a docking experiment [40, 41]. But the default exhaustiveness value is 8 and the creators claim that it should increase the time linearly and decrease the probability of not finding the minimum exponentially [40, 41], hence increasing this value leads to an exponential increase in the probability of finding the energy minima.

The whole protein conformational space was searched, using grid box dimensions 60×62× 48 A°. Following exhaustiveness values were tested in this study: 8, 16, 32, 64, 128, 256, 512, 1024, 2048 and 4096.

Principal component (PC) [42] and cluster analysis using K-means algorithm [43] was performed (Additional file 1: Figures S4, S5). The results demonstrated that the number of interaction sites doesn’t change in the interval using exhaustiveness from 1024 to 4096. Exhaustiveness value of 1024 was chosen as it provides good results, good speed and thorough sampling of the docked configurations. Exhaustiveness value was increased to a value of 1024 and a maximum number of binding modes to generate set to 20. After that 100 independent docking calculations were carried out with random initial seeds.

We also used other docking programs for comparison using rDock [44] and FlexAid [45], We performed 10 docking simulation for each program and generated 20 docked poses.

### Analysis of docking conformations and trajectories

We performed the analysis of docking conformations and trajectories by a combination of Autodock Vina [40], Gromacs [32] in addition to custom python scripts, which uses pandas [46], scikit-learn [47] and MDTraj [48].

Here is the list of program packages we used for analysis:

- The MDTraj python library [48] was used for the trajectory analysis and LigPlot+ for 2D diagrams of 3-0-C12-HSL-LasR interactions [49],
- Plot visualization was done using matplotlib [50] and Seaborn package [51].
- Figures and videos were prepared with PyMOL [52], VMD [53] and UCSF Chimera [54].

The analysis protocol approach was similar to the works of Wolf et al. [55] and Zamora et al. [56] and the following techniques were used for the analysis:

- Principal component analysis (PCA) is an unsupervised statistical technique that is often used to make data easy to explore and visualize. PCA tries to explain the maximum amount of variance with the fewest number of principal components [42], The process of applying PCA to a protein trajectory is called Essential Dynamics (ED) [56-58]. PC analysis, performed on Cartesian coordinates has become an important tool for the examination of conformational changes.
- Cluster analysis is another unsupervised technique that tries to identify structures within the data. It is a data exploration tool for dividing a multivariate dataset into groups. Clustering algorithms can be grouped into partitional and hierarchical clustering approaches [59].

### NMR calculations

Finally, the trajectory of LasR monomer simulation was used for the calculation of chemical shifts. Sparta+ [60] and ShiftX2 [61] were used to predict the chemical shifts of backbone atoms of the protein with the help of wrapper functions of MDTraj [48].

### Binding energy calculation

g_mmpbsa [62] was used for the evaluation of binding energies. It was also used for the estimation of the energy contribution of each residue to the binding energy, g_mmpbsa calculates the relative binding energy rather than the absolute binding energy, so the entropy contribution (TΔS) is not evaluated [62], We cut out from the 300ns simulations production trajectories of 10ns with the interval of every 120 ps for the calculation of all energy components of binding energies.

### Sequence Conservation

To find out whether 3-0-C12-HSLs interact with conservative amino acid residues we performed multiple sequence alignment (MSA) using msa package [63]. ClustalW [64], Clustal Omega [65] and Muscle [66] within msa package were used for multiple sequence alignments. We performed multiple sequence alignment of the LasR protein between the closely related species of *P. Aeruginosa.*

The following sequences were used for sequence alignment:

- WP_054058449.1 LuxR family transcriptional regulator [*Pseudomonas fuscovaginae*]
- NP_250121 transcriptional regulator LasR [*Pseudomonas aeruginosa* PAOl]
- KFC75736.1 LuxR-type transcriptional regulator [Massilia sp. LC238]
- WP_018433960 LuxR family transcriptional regulator [Burkholderia sp. JPY251]
- WP_042326260.1 LuxR family transcriptional regulator [Paraburkholderia ginsengisoli]
- WP_012426170.1 LuxR family transcriptional regulator [Paraburkholderia phytofirmans]
- WP_027776298.1 LuxR family transcriptional regulator [Paraburkholderia caledonica]
- WP_027214716.1 LuxR family transcriptional regulator [Burkholderia sp. WSM2232]
- WP_041729325.1 LuxR family transcriptional regulator [Burkholderia sp. CCGE1003]
- CBI71275.1 UnaR protein [Paraburkholderia unamae]
- CAP91064.1 BraR protein [Paraburkholderia kururiensis]
- WP_003082999.1 transcriptional activator protein LasR [*Pseudomonas*]
- WP_012076422.1 MULTISPECIES: LuxR family transcriptional regulator [*Pseudomonas*]
- WP_050376898.1 MULTISPECIES: LuxR family transcriptional regulator [*Pseudomonas*]
- WP_050395760.1 LuxR family transcriptional regulator [Pseudomonas aeruginosa]

### Protein-Protein Docking

When docking homology models, it is best if there is an experimental evidence to suggest the general interaction site (within ~10 Å). Representative structures from molecular dynamics simulations were used for protein-protein docking using ClusPro [67], From the experimental X-ray data, it was found that ‘Trpl52’, ‘Lys 155’ and ‘Asp 156’ play an important role during dimerization.

The distances between ‘Trpl52’ from chain A and ‘Aspl56’ from chain B was 0.276 nm, ‘Aspl56’ from chain A and ‘Lysl55’ from chain B of the crystallographic structure was 0.279nm. These residues were used as attraction constraints.

### Entire flowchart

The whole methodology is presented as a flowchart for a better comprehension:

- LasR protein sequence taken from UniprotKB (sequence ID P25084) and performed blastp of LasR to identify suitable templates for reconstruction.
- HH-pred web server was used for the reconstruction of LasR monomer structure and validated using Verify 3D, Procheck, Prosa.
- The 3D model of 3-0-C12-HSL acquired from PubChem web server.
- Docking of 3-0-C12-HSL ligand with the LasR monomer performed using Autodock Vina, later we corroborated with rDock, FlexAid.
- PCA and cluster analysis performed on docking conformations.
- Extraction of centroid conformations from cluster analysis.
- Ligand parameters generated using Acpype interface in the framework of the AMBER force field.
- Using centroid conformations as starting points for molecular dynamics simulations using Gromacs.
- Analysis of molecular dynamics trajectory files using MDTraj.
- Relative binding energy calculation and contribution of residues to binding using g_mmpbsa.
- Sequence alignment using msa package with ClustalW, Clustal Omega and Muscle algorithms.
- Performed protein-protein docking using ClusPro.

## Results and discussion

### Conformational changes of LasR without 3-0-C12-HSL

We performed a simulation run of 100 ns using a standard MD protocol for the assessment of conformational changes of LasR monomer (Additional file 2). The overall stability of the molecule was assessed using the root-mean-square-deviation (RMSD) of the protein atoms. RMSD was calculated with reference to the initial frame through the time of MD run (Additional file 1: Figure S6). Another suitable measurement for the stability of the LasR monomer structure is the radius of gyration (Additional file 1: Figure S7).

During the examination of MD trajectories. Principal Component (PC) analysis [42] is usually used for the visualization of the motions of the system (Figure 2). Generally to capture more than 70% of the variance in the trajectory data the first handful of components are sufficient [56], PC analysis can uncover the fundamental movements contained in an MD trajectory, however, it does not group the snapshots into different clusters [59], This can be accomplished by the clusterization of the PC data.

**Fig. 2.**
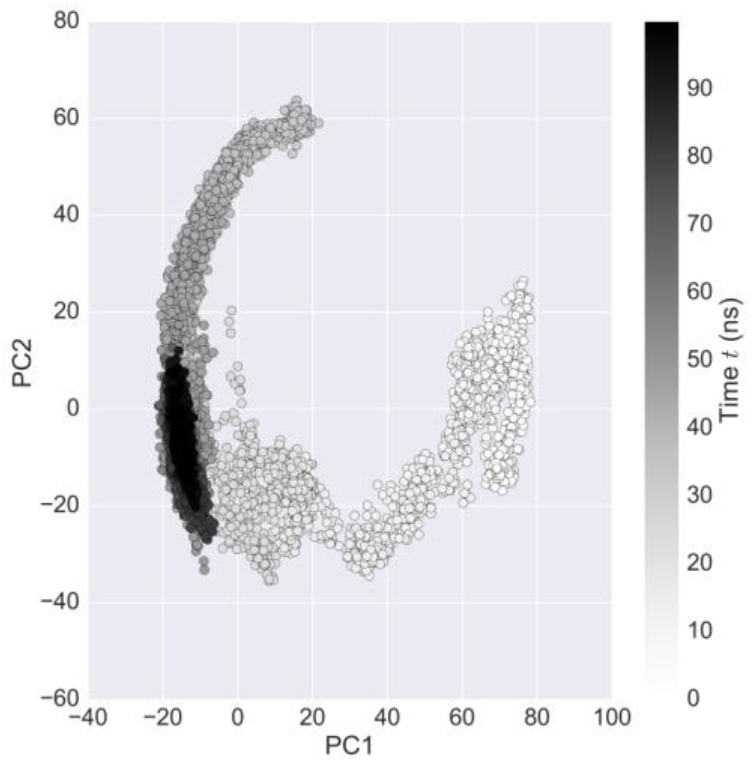
LasR monomer simulation trajectory projected onto its first two principal components. The black scale indicates the time progression from 0 ns (white) to 100 ns (black).

For the selection of initial starting structure of the LasR for docking study, we performed cluster analysis on the LasR monomer run for the selection of a starting point. By identifying a distinct representative structure of the most populated cluster, this will allow performing blind docking on the whole structure. It should be also noted that cluster analysis allows evaluating the frequent conformations of LasR. For the clustering analysis, we chose the Agglomerative Clustering algorithm with Ward linkage [68] as the most appropriate as they are deterministic, allowing for reproducibility of resulting clusters, thus minimizing the amount of bias.

We performed several rounds of Agglomerative clustering with Ward linkage (details can be found in the methods section). The accuracy of the clustering was assessed by the help of the Davies-Bouldin Index (DBI) [69], Dunn Index [70], Silhouette Score [71], and the pseudo-F statistic (pSF or Calinski Harabasz) [72] metrics (Additional file 1: Figure S8). An optimal number of clusters were chosen, simultaneously accounting for a low DBI, high Silhouette, high Dunn Index and high pSF values.

The distribution of clusters over simulation is visualized in Figure 3 and the four cases are cluster 4 (black) at the beginning of the simulation (after equilibration), cluster 2 (green) in the middle, cluster 1 (light green) and cluster 3 (dark blue) at its end.

**Fig. 3.**
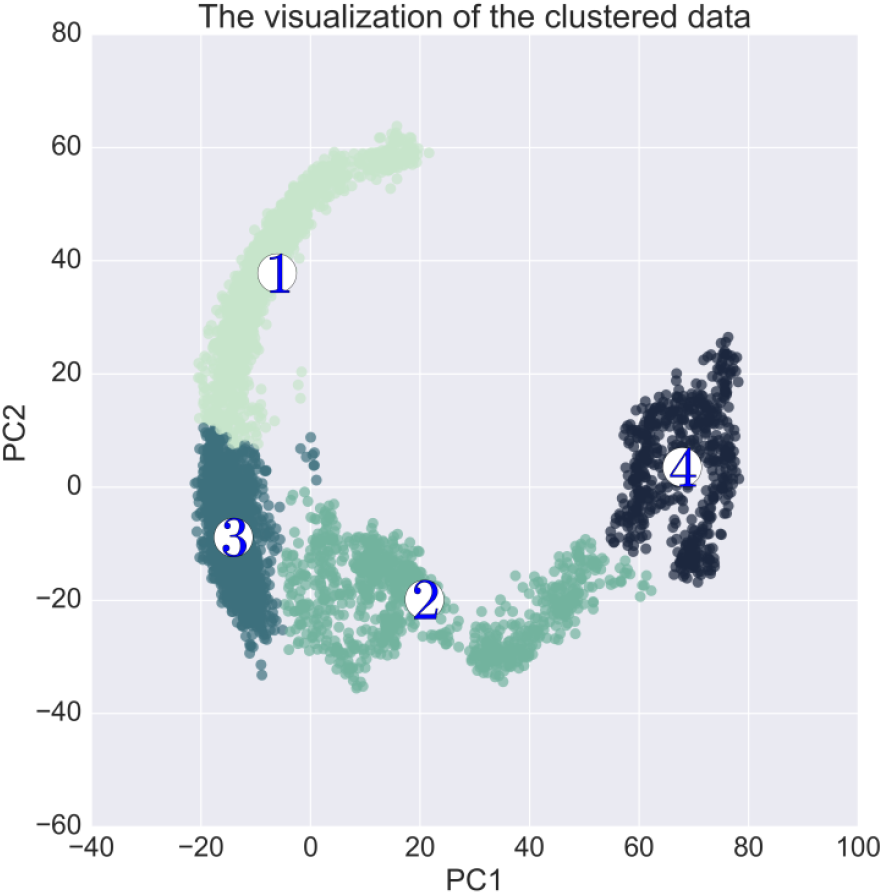
Clustering results of ward-linkage algorithm formed by first two PCs. The entire MD trajectory data was used for the clusterization.

The clusterization defined by the first two PCs (Figure 4) provides a coherent picture and it also supported by a good DBI, Dunn, Silhouette Score and pSF value (Additional file 1: Figure S8). Figure 4 shows clearly that the simulation has converged.

**Fig. 4.**
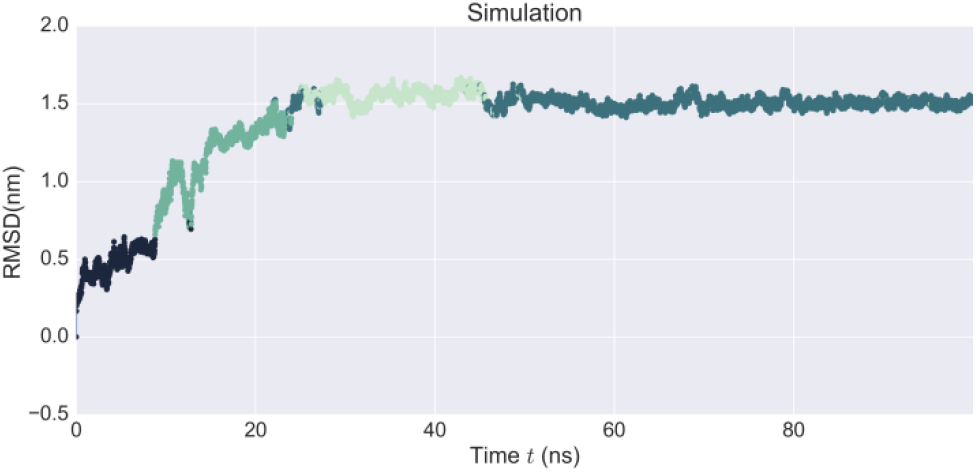
Colour-coded RMSD of the simulation obtained from Ward-linkage cluster analysis. The colours represent the clusters that are demonstrated in Figure 4.

For the validation of the quality of molecular dynamic simulation, theoretical NMR shifts were calculated using Sparta+ [60] and ShiftX2 [61]. For the NMR shifts calculation, snapshots from cluster 3 (Figure 3 and Figure 4) were used.

In Figure 5 you can see that there is a strong linear relationship between experimental and simulated ShiftX2 NMR shift values (Additional file 1: Figure S9), thus this demonstrates the quality of MD simulation.

**Fig. 5.**
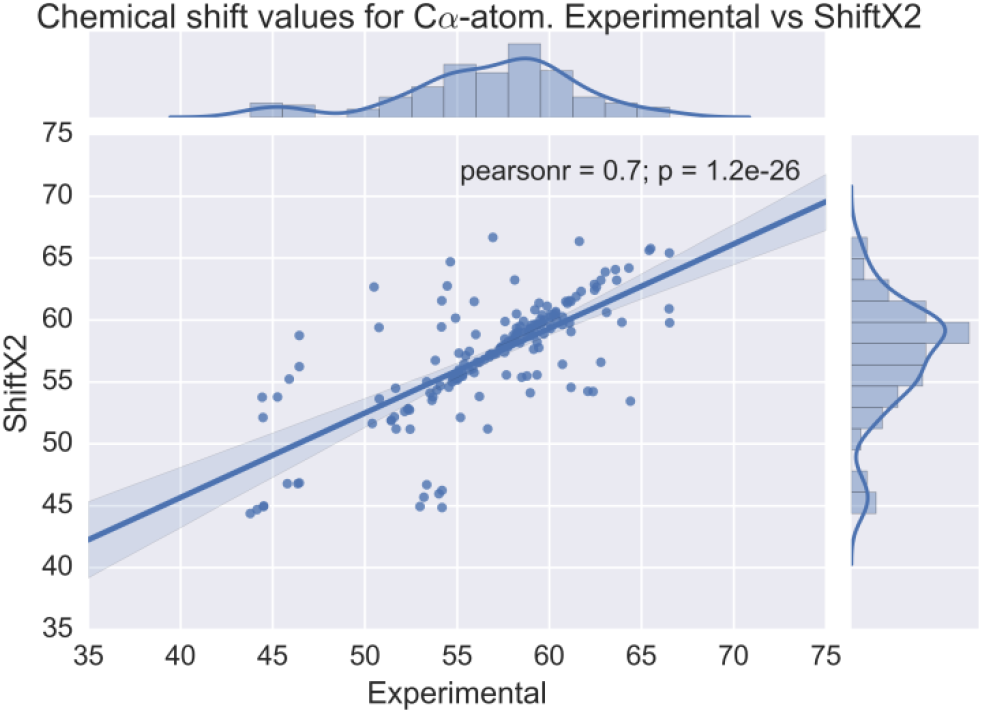
Chemical shifts of Cα-atoms. Experimental vs ShiftX2 predictions.

Figure 6 shows that there is a strong linear relationship as well between experimental and simulated Sparta+ NMR shift values (Additional file 1: Figure S9).

**Fig. 6.**
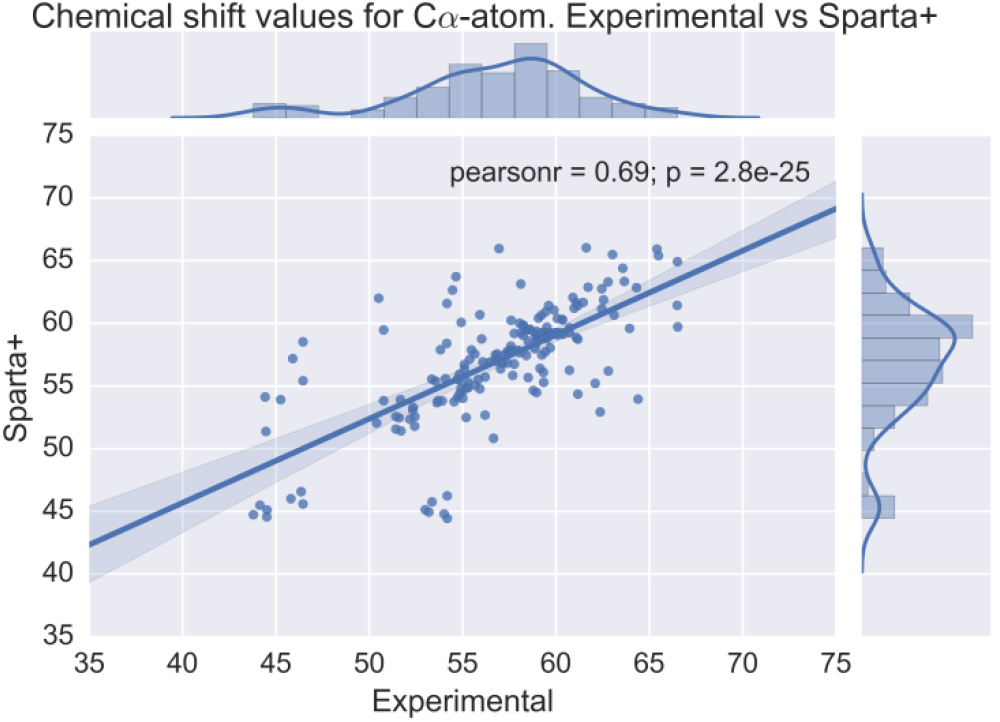
Chemical shifts of Cα-atoms. Experimental vs Sparta+ predictions.

After that, for docking study, the representative structure (Figure 7c) was extracted from cluster 3 (Figure 3 and Figure 4).

**Fig. 7.**
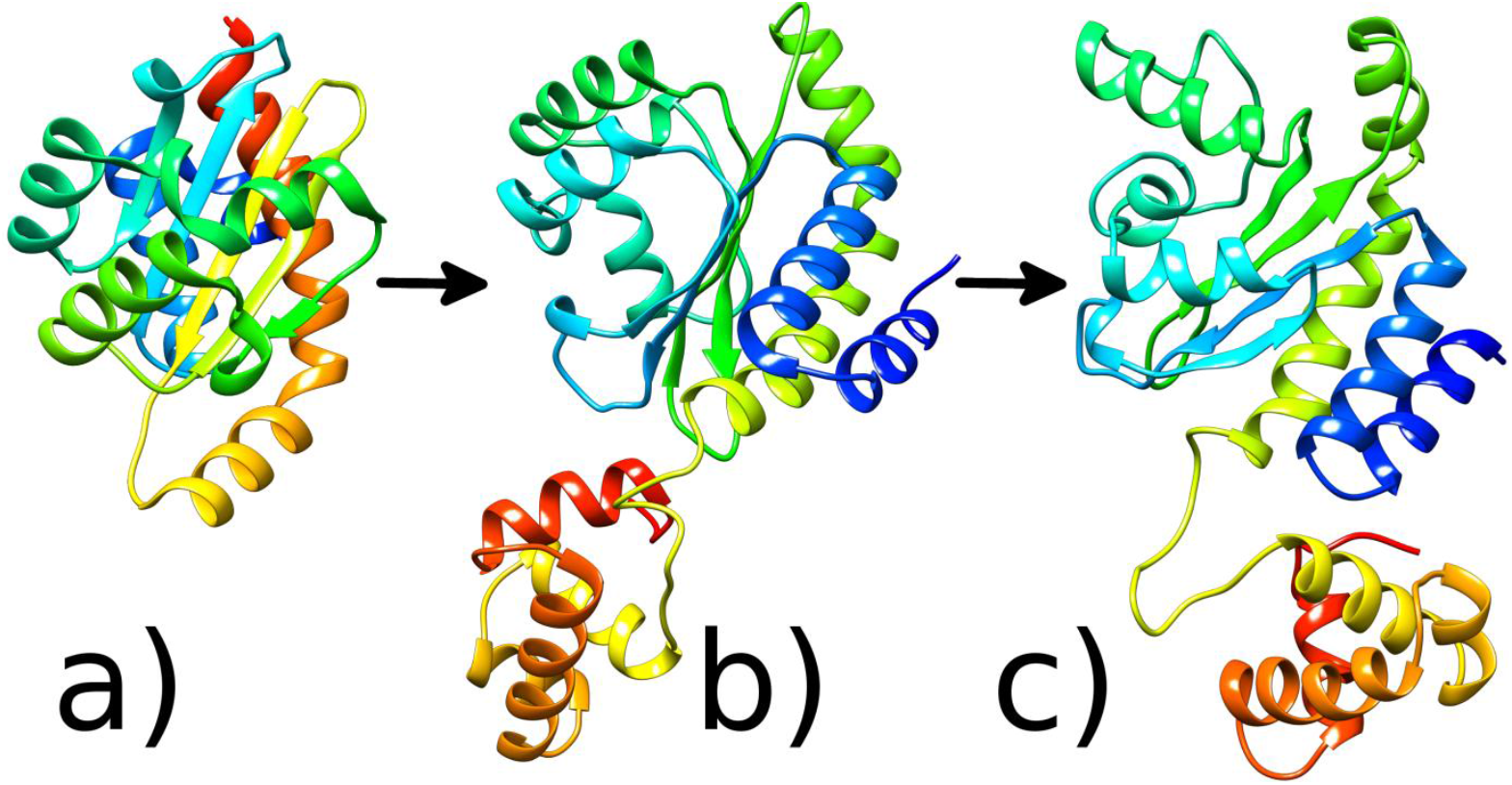
3D LasR protein structures: a) Crystal structure of the N-terminal LBD of LasR protein b) Homology Modelling c) Representative Structure after 100ns MD run. Images generated with UCSF Chimera [49].

The representative structure of the LasR system and its differences from the homology modelled structure are highlighted in Figure 8c. Secondary structure analysis of the representative structure was performed as well using PDBSum [73] (Additional file 1: Figure S10). To our opinion, this is a complete study to include the dynamics of the full-length LasR molecule (residues 1 to 239). It has also been shown that the dynamics of the complete C-terminal region of LasR modulate N-terminal region, as will be discussed later.

### Docking analysis of 3-0-C12-HSL with LasR monomer

In this study, molecular docking approach was used to inspect the possible binding modes of 3-0-C12-HSL in LasR monomer. PCA and cluster analysis were performed on docking data (Figure 8). The results show three binding sites, cluster 2 corresponds to experimental data, while cluster 1 and cluster 3 do not (Figure 8). These results clearly support the findings of Bottomley et al. [18], where it was also demonstrated that 3-0-C12-HSL binds to LBD.

We performed several rounds of K-Means clustering (details are available in the section of methods). The accuracy of the cluster analysis was evaluated using the DBI [69], Dunn Index [70], Silhouette score [71] and the pSF [72] metrics (Additional file 1: Figure Sll). An optimal number of clusters were chosen for docking results, simultaneously accounting for a low DBI, high Dunn, high Silhouette and high pSF values.

We generated 2000 docked poses and performed representative structure extraction for use in MD simulations of the LasR 3-0-C12-HSL binding sites. The resulting representative structures from each cluster are shown in Figure 9. These cluster representative structures were produced by finding the centroid conformations.

**Fig. 8.**
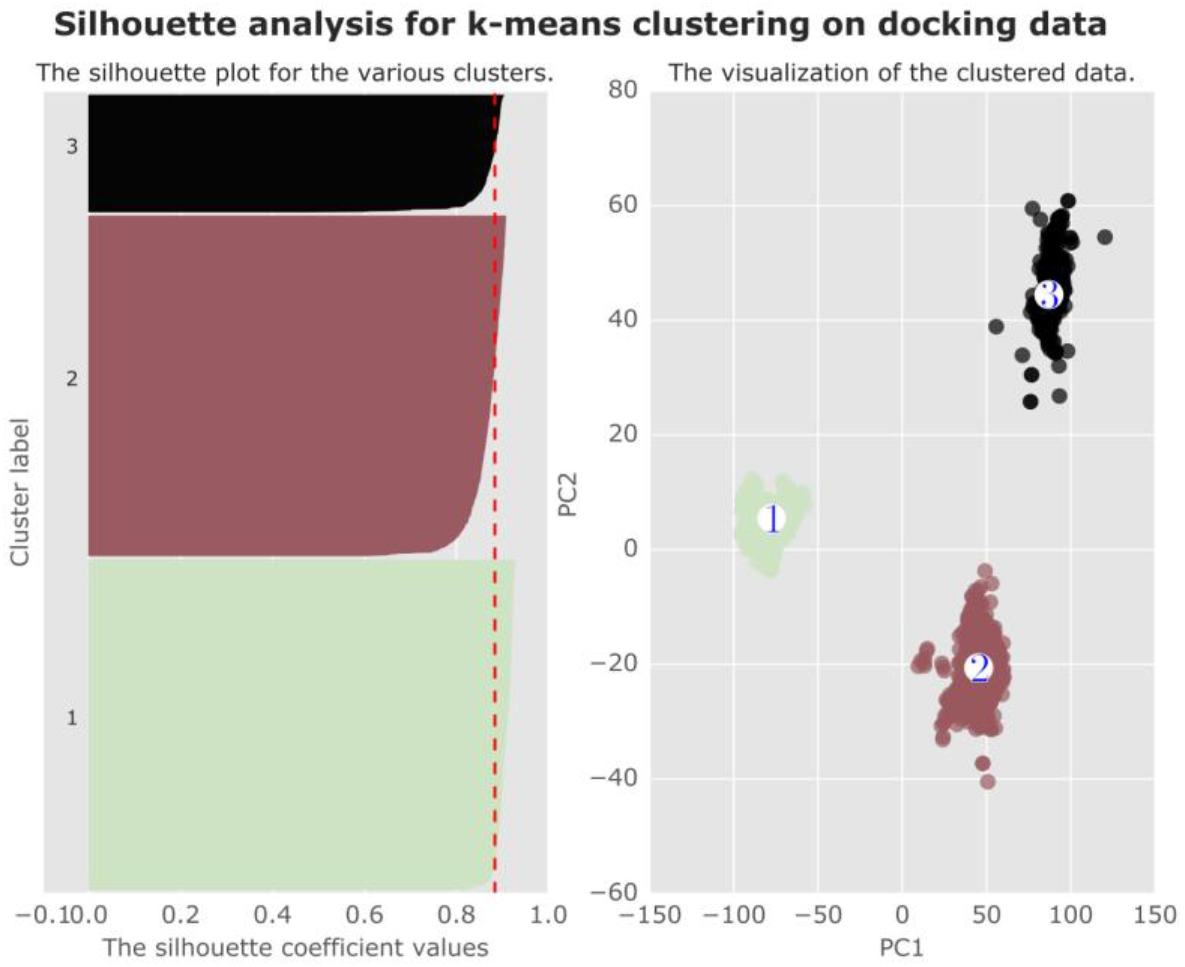
Silhouette plot (left) and clustering results (right) on the docking data using k-means algorithm formed by first two PCs.

Representative structures from each cluster were extracted. The binding energy for the representative structure of cluster 1 is -5.4 kcal/mol, as for the mean binding affinity for the whole cluster is -5.257 ±0.233 kcal/mol (Additional file 1: Figure S12). Cluster 1 contains 839 docked poses from 2000, about 41.95%. For cluster 2 the binding affinity for the representative structure is -5.1 kcal/mol and for the whole cluster -5.593 ±0.386 kcal/mol (Additional file 1: Figure SI3) and this one corresponds to the experimental binding site data [18], Cluster 2 contains 864 docked poses from 2000, about 43.2%. For cluster 3, the representative structure features the highest binding affinity -5.7 kcal/mol and for the whole cluster -5.264 ±0.27 kcal/mol (Additional file 1: Figure SI4). Cluster 3 contains 297 docked poses from 2000, about 14.85%, which is a rather unstudied area.

There is a huge amount of literature that suggests that molecular docking is not appropriate for the prediction of binding affinity or binding poses of protein-ligand complexes, however, they can still provide important information [39, 74],

**Fig. 9.**
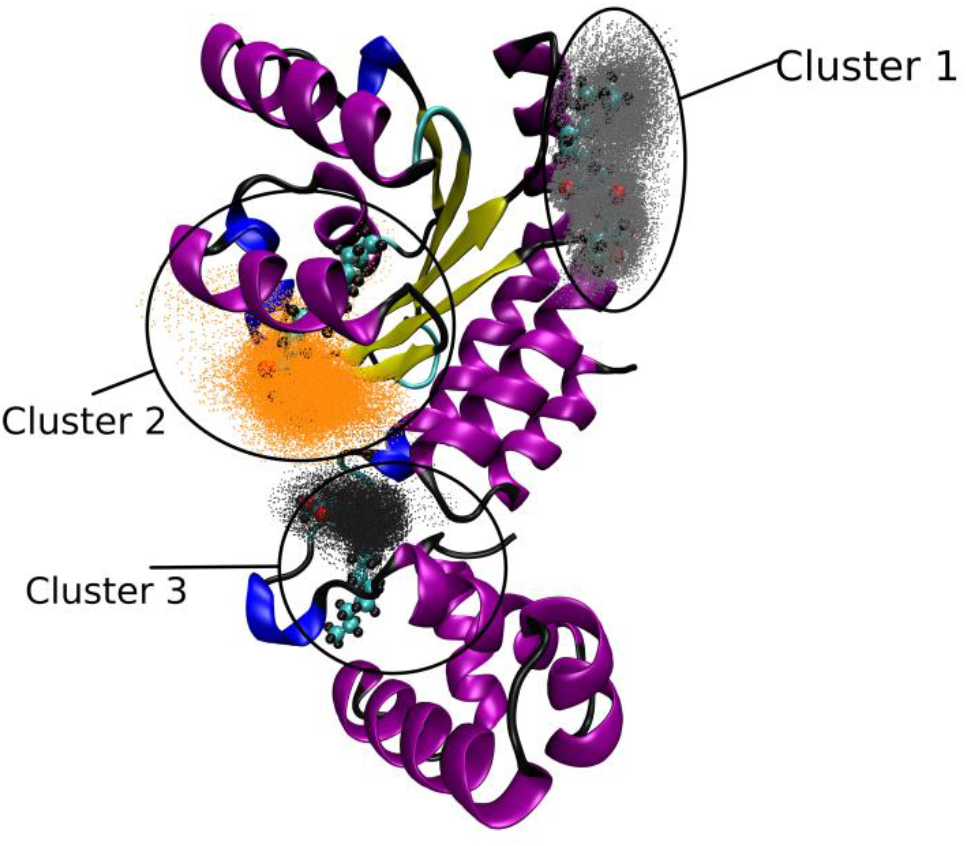
3D visualization of the analysed docking data with their representative structures and clusters.

Later we corroborated the blind docking with other molecular docking software as well, which includes Autodock Vina [40], rDock [44] and FlexAid [45] (Figure 10). PC analysis of the various docking programs was performed for easier comprehension (Additional file 1: Figure SI 5).

**Fig. 10.**
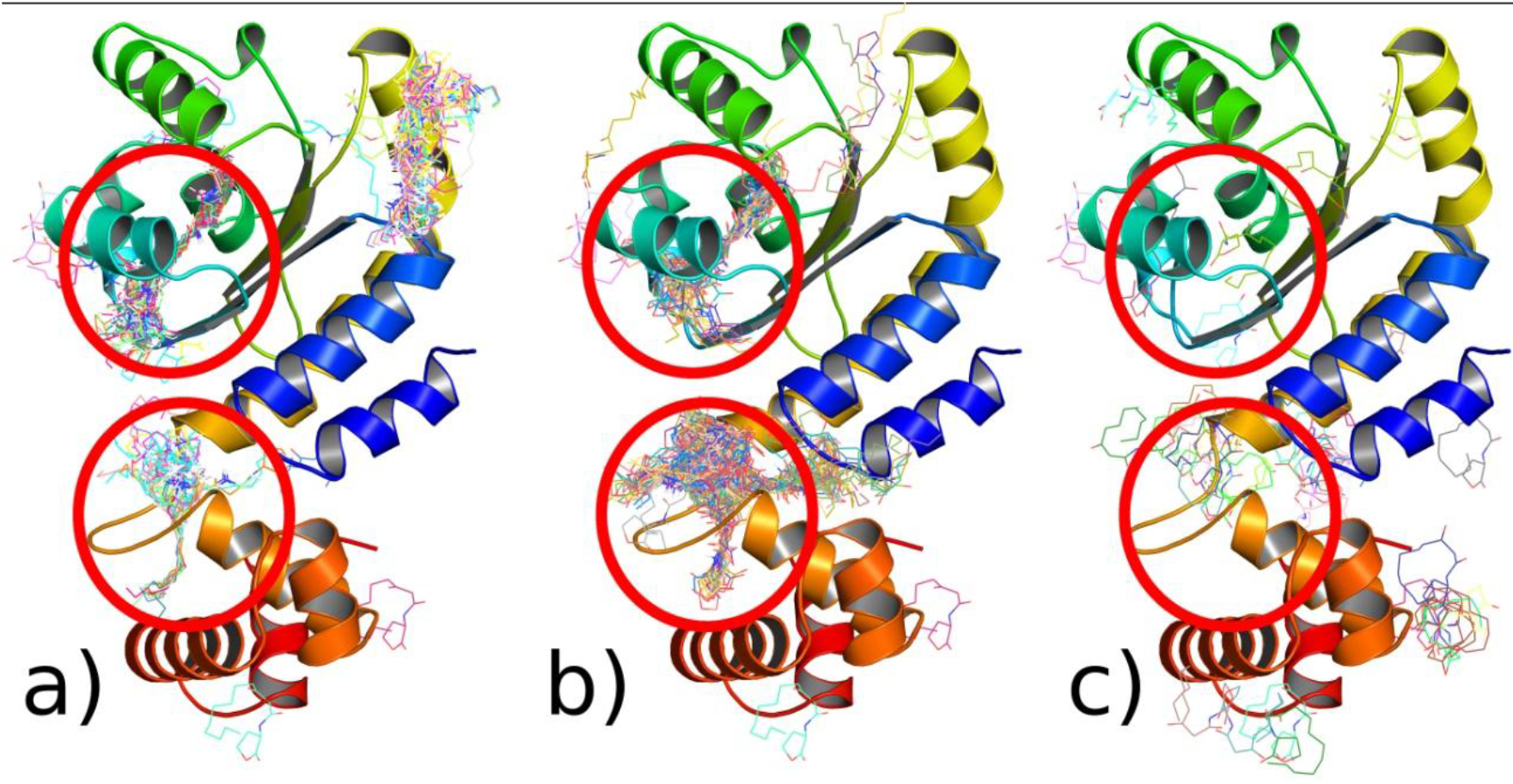
Blind docking with various molecular docking programs. Red circles - binding interactions that are characteristic to all programs a) Autodock Vina b) rDock c) FlexAid.

### Binding Modes of 3-0-C12-HSL

We performed three 300 ns simulations using standard MD protocol. Overall, 900ns of simulation data was used for the analysis of 3-0-C12-HSL interaction with LasR monomer. The representative structures were taken from docking results (Figure 9) and used as starting points for MD simulation with LasR. Simulations were conducted for sufficient time to allow the positions of 3-0-C12-HSL to reach equilibrium in LasR molecule.

The overall stability of the molecule was assessed using the mass-weighted RMSD of the backbone atoms. RMSD was calculated with reference to the initial snapshot for the different independent MD runs. Figure 11 shows that Simulation 2 and 3 experience a substantial RMSD deviation from the initial starting point. Simulation 2 corresponds to cluster 2 in docking simulations, while Simulation 3 corresponds to cluster 3 (Figure 11). Simulation 1 which corresponds to cluster 1, the molecule of 3-0-C12-HSL did not fixate and reach equilibrium, so further research was not performed (Figure 11) (Additional file 3). Simulation 2 (Additional file 4) shows that after 230ns, the structure becomes stable. While Simulation 3 (Additional file 5) changes its conformation in 100ns and becomes stable between 260ns and 300ns.

**Fig. 11.**
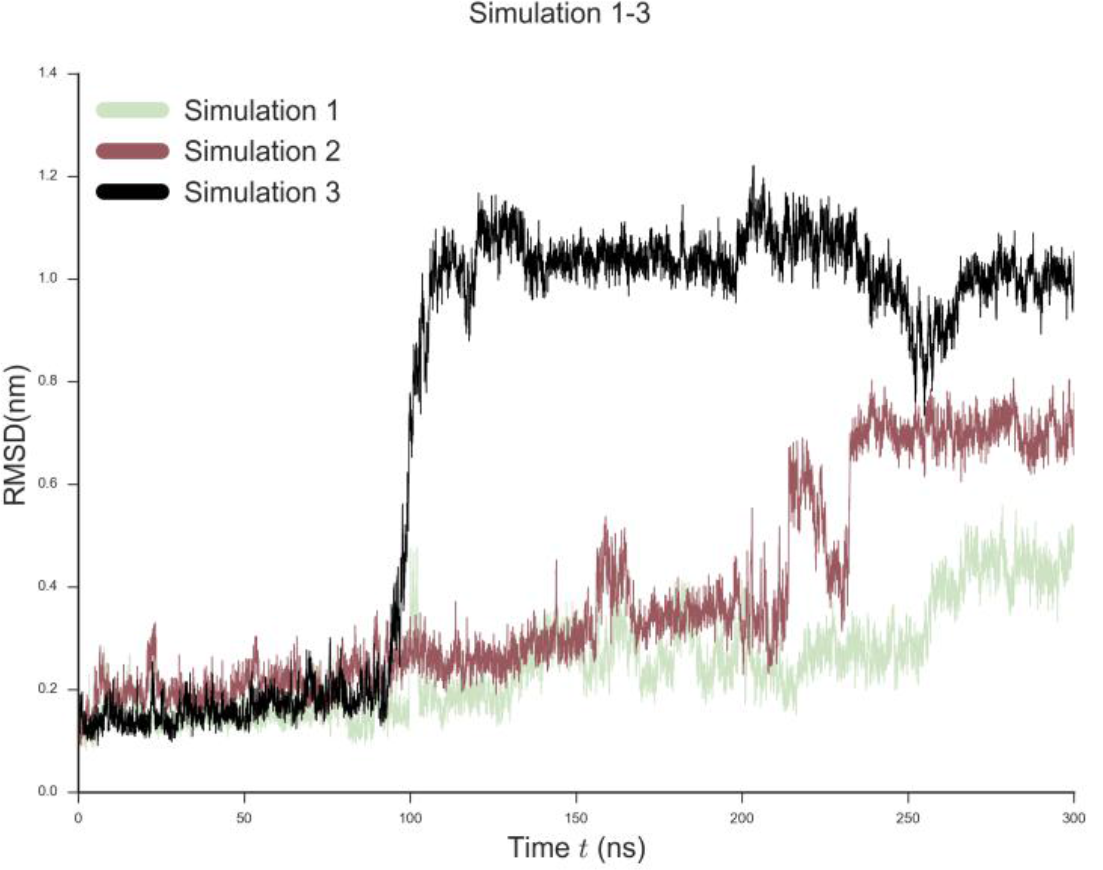
RMSD evolution during 300 ns of simulation for 3 independent runs using representative structures of 3-0-C12-HSL docking poses as starting points.

The root-mean-square fluctuation (RMSF) was used for the assessment of the flexibility of the LasR monomer. In Figure 12 the average per-residue RMSF for each cluster simulation runs is plotted.

**Fig. 12.**
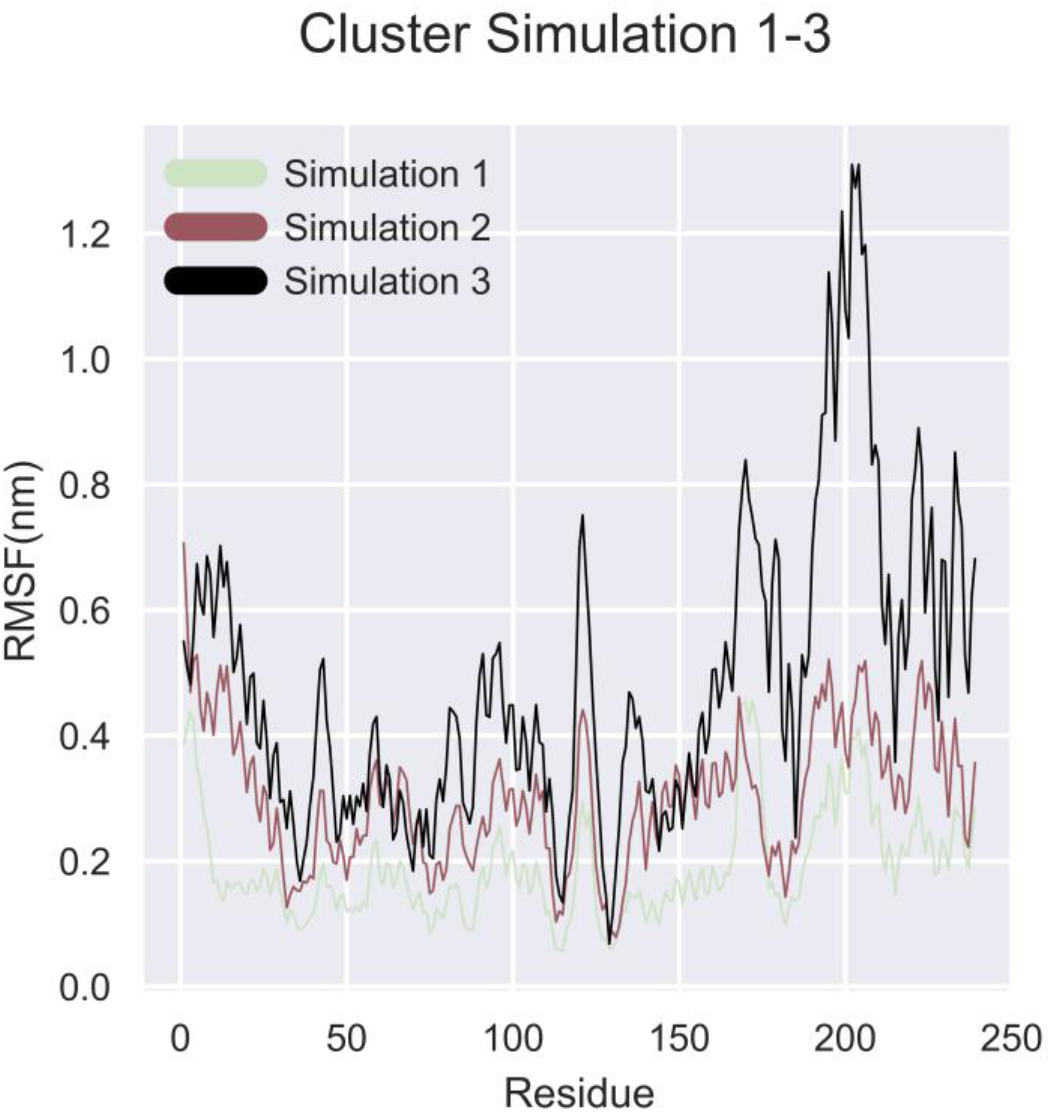
RMSF of Cα-atoms of LasR for each individual run.

From Figure 12 it visible that the residues from 165 to 176, which correspond to beta turns in the SLR of LasR, are of high mobility (Additional file 1: Figure S10). Simulations 2 and 3 show that 3-0-C12-HSL has two binding modes, one with LBD, which corresponds to experimental data and simulation 3 with the LBD-SLR-DBD bridge of LasR.

PCA and Cluster analysis were performed on simulation 2 (Additional file 1: Figure SI6) as well as hydrogen bond analysis based on cutoffs distance and angle according to the criterion of Wernet-Nilson [75] using MDTraj [48]. Over the course of cluster 1, 3-0-C12-HSL with LasR established an average of 0.655±0.651 of hydrogen bonds, while in cluster 2 the average is 0. 042±0.202 (Additional file 1: Figure S16). Over the course of simulation 2, 3-0-C12-HSL establishes a large number of hydrophobic contacts with amino acid side chains of the LBD of LasR protein (Additional file 1: Figure SI8), a phenomenon that is not unexpected, given the large hydrophobic surface area of the LasR LBD and the low solubility of 3-0-C12-HSL in water. In simulation 2, 3-0-C12-HSL has hydrophobic interactions mainly with amino acids from H3, H5 and SI (Additional file 6, 7, 8). RMSD analysis of the conformations between LasR with/without 3-0-C12-HSL in LBD is equal to 7.027 Å (Figure 13).

**Fig. 13.**
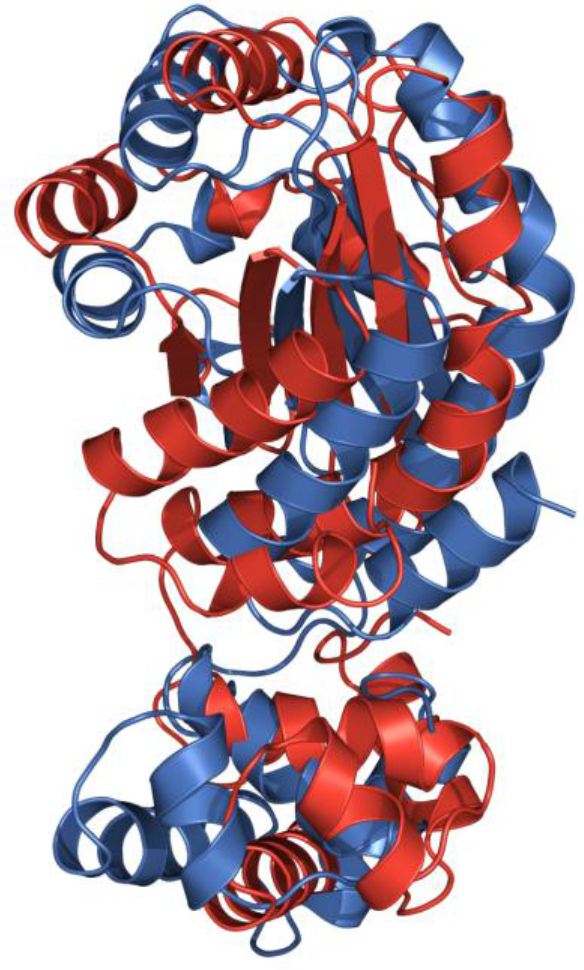
LasR monomer structures: without 3-0-C12-HSL (blue colour), with 3-0-C12-HSL bound to LBD (red colour).

Residues that participate in hydrophobic interactions are shown in Figure 14 and Additional file 1: Figure SI8.

**Fig. 14.**
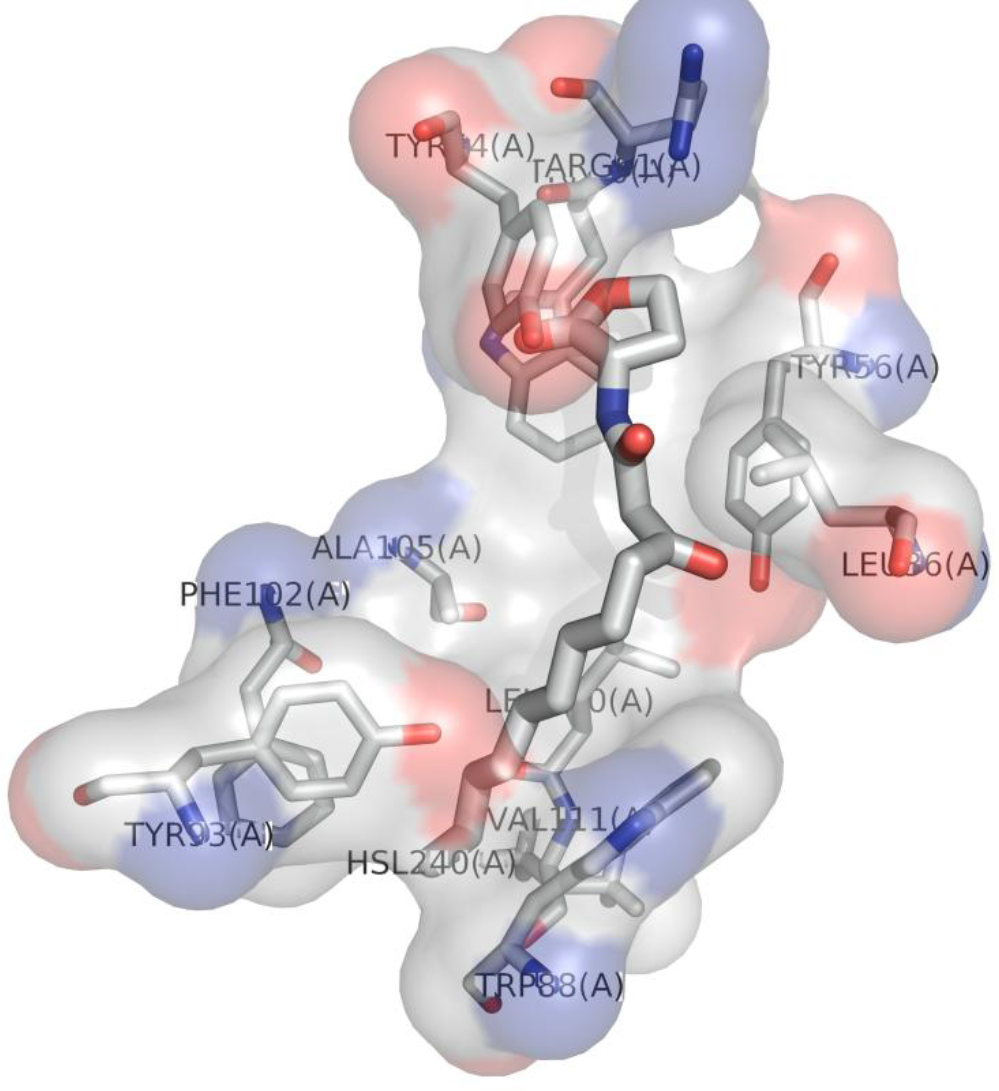
Insertion of 3-0-C12-HSL in LBD of LasR, from centroid snapshot of cluster 2 in simulation 2 (Additional file 1: Figure SI6).

The second binding mode involves the interaction of 3-0-C12-HSL with the LBD-SLR-DBD bridge. PCA, cluster, and hydrogen bond analysis were also performed on simulation 3 (Additional file 1: Figure S19). Over the course of cluster 2, 3-0-C12-HSL with LasR established an average of 1.506±0.742 hydrogen bonds, for cluster 3 the average 0.228±0.492 (Additional file 1: Figure S19), while in cluster 1 the average is 0.652±0.654. Over the course of simulation 3, 3-0-C12-HSL establishes hydrogen bonds and a large number of hydrophobic contacts with amino acid side chains in the beta turns in the SLR of LasR protein (Additional file 1: Figure S21). In simulation 3, 3-0-C12-HSL forms hydrogen bonds mainly with ‘Lysl82’ and ‘Lcul77’ of the beta turns in the SLR of LasR (Additional file 1: Figure S21). RMSD analysis of the conformations between LasR monomer and LasR bound to 3-0-C12-HSL in DBD is equal to 1.677 A (Figure 15).

**Fig. 15.**
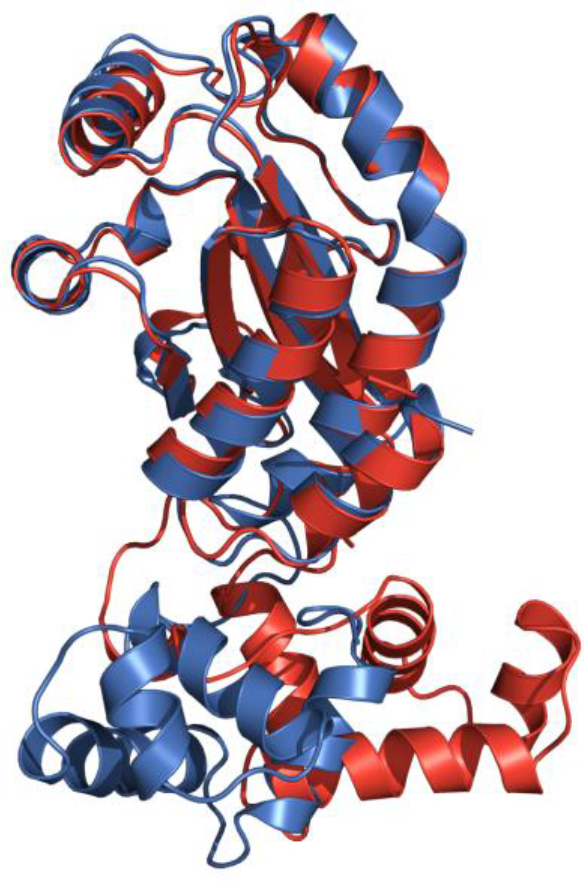
LasR monomer structures: without 3-0-C12-HSL (blue colour), with 3-0-C12-HSL bound to LBD-SLR-DBD bridge (red colour).

Residues that participate in hydrogen and hydrophobic interactions are shown in Figure 16.

**Fig. 16.**
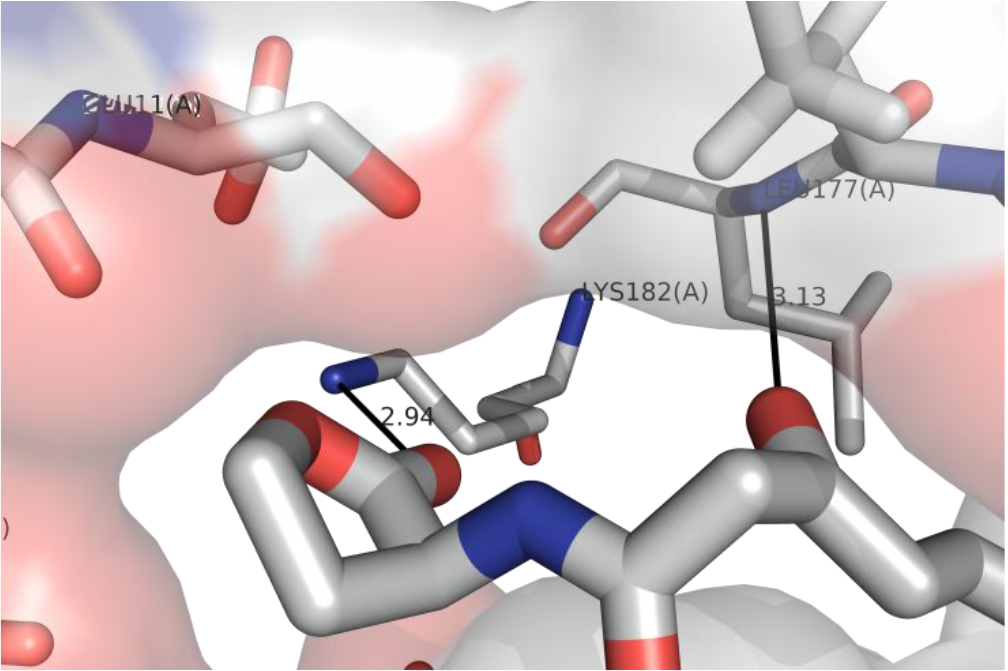
Putative hydrogen bonds of 3-0-C12-HSL with Lysl82 and Leul77 from centroid snapshot of cluster 1 of simulation 3 (Additional file 1: Figure SI 9).

Complete data sets for MD simulations are available as Supporting Information (Additional file 1: Figures S16-S21).

### Binding Energy of 3-0-C12 to LasR and Sequence Conservation

In order to analyse the binding sites in detail, MMPBSA [62] binding energy calculation was performed for each binding site based on the trajectories. The binding energy calculation demonstrated a very interesting results and that is that the interaction with LBD-SLR-DBD bridge is higher than with the LBD (Table 2). More detailed analysis on the energy terms showed that Van der Waals, electrostatic interactions and non-polar solvation energy contribute negatively to the binding energy while polar solvation energy contributes positively. And van der Waals interaction contributes most in the terms of negative contribution for both cases, but for the interaction with the LBD-SLR-DBD bridge or “the bridge” the electrostatic interaction is 3.6 higher than with the LBD interaction.

**Table 2.**
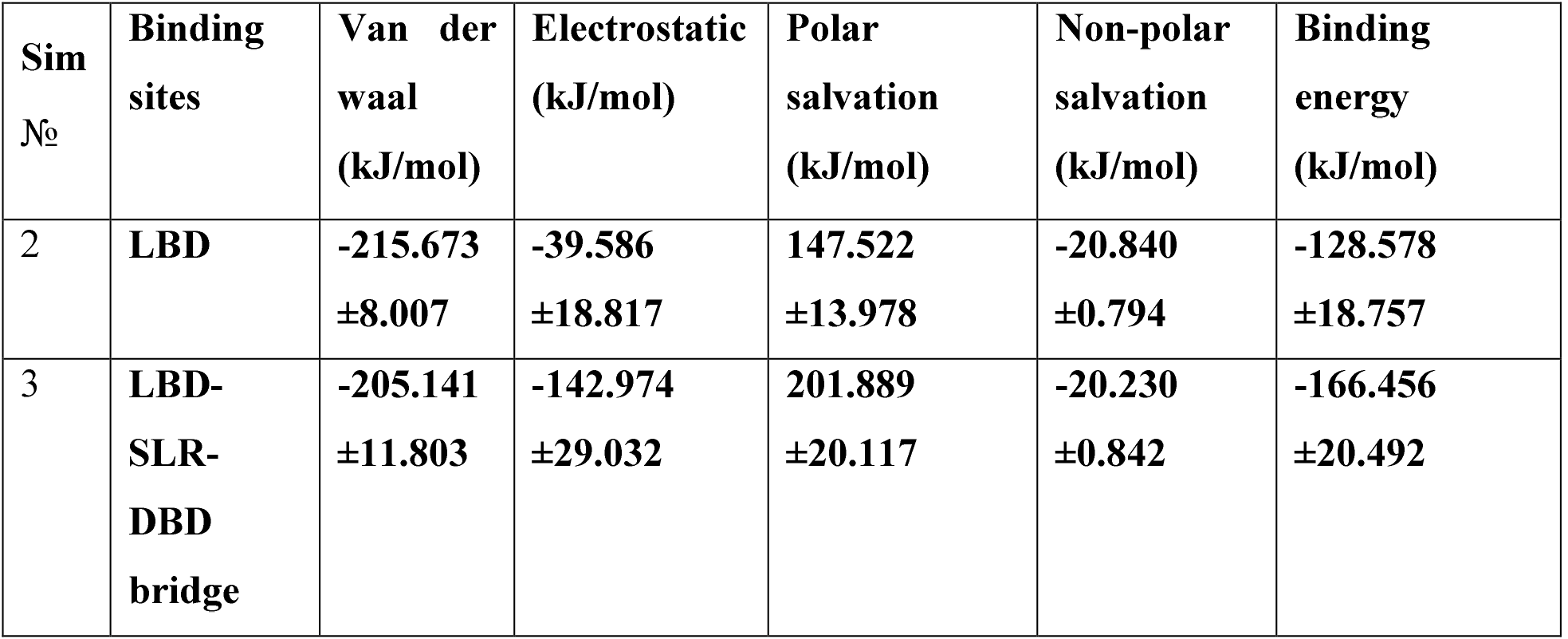
Relative Binding energy using g_mmpbs on simulation data.

This suggests that van der Waals, electrostatic interaction and non-polar solvation energy together contribute to the stability of LasR-3-0-C12-HSL binding complex. To asses with which conserved residues in But to find out we needed to find out which amino acid residues were involved with the interaction. We performed analysis of Energy contribution of residues to binding for both simulations obtained from MMPBSA calculation using g_mmpbsa. Sequence alignment was performed using msa package [63], ClustalW [64], Clustal Omega[65] and Muscle[67] algorithms were used for sequence alignment. Full alignments using the aforementioned algorithms are available as supplementary materials (Additional file 6, 7, 8). The binding state with LBD with binding energy ~-128.578 kJ/mol (Table 2), which the residues Tyr64, Tyr56. Trp88. Leu36 and Trp60 contribute most to (Figure 17).

**Fig. 17.**
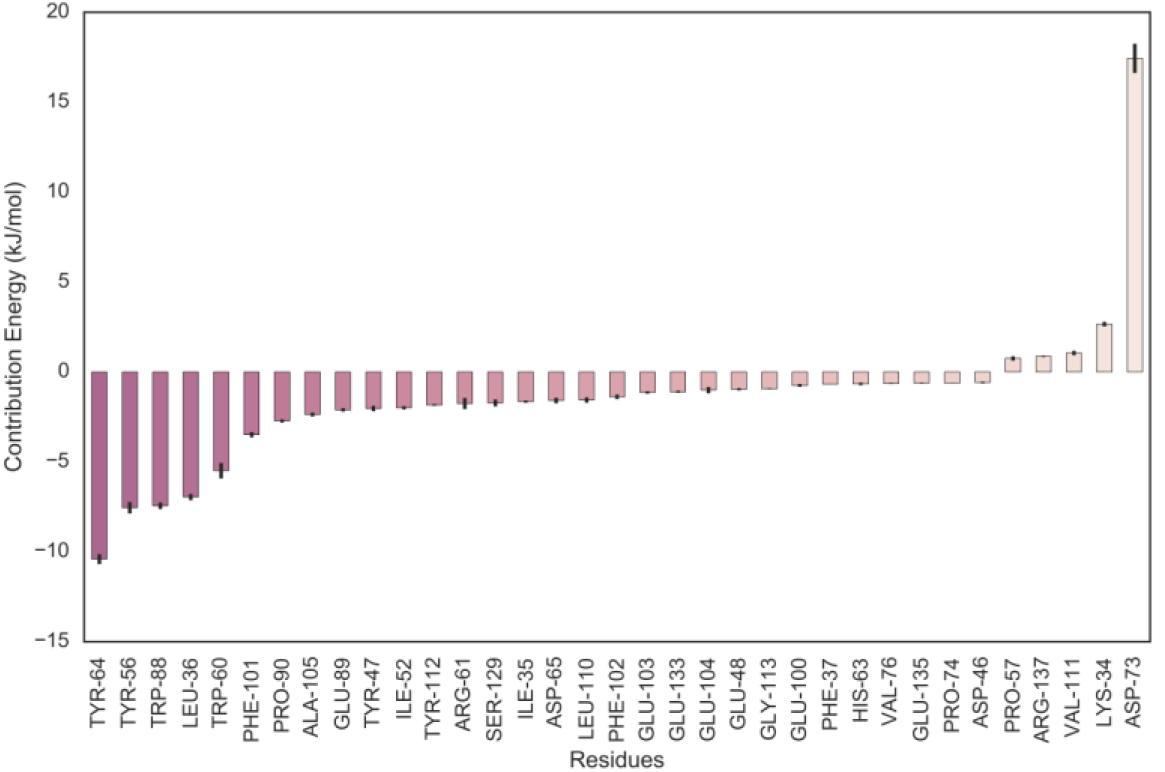
Energy contribution of LBD amino acid residues from simulation 2.

Residue interaction and sequence alignment show that 3-0-C12-HSL interacts with very conservative amino acid that includes Trp60 which corresponds to experimental data [76], It also involves other conserved amino acids such as Tyr64, Asp73, Pro74. Val76. Phel 02. Phel 03. Alai05. and Glyll3 (Fig. 18). 3-0-C12-HSL interacts with 10 fully conserved amino acids. In total 16 amino acids participate in the interaction, where amino acid conservation is more than 75%.

**Fig. 18.**
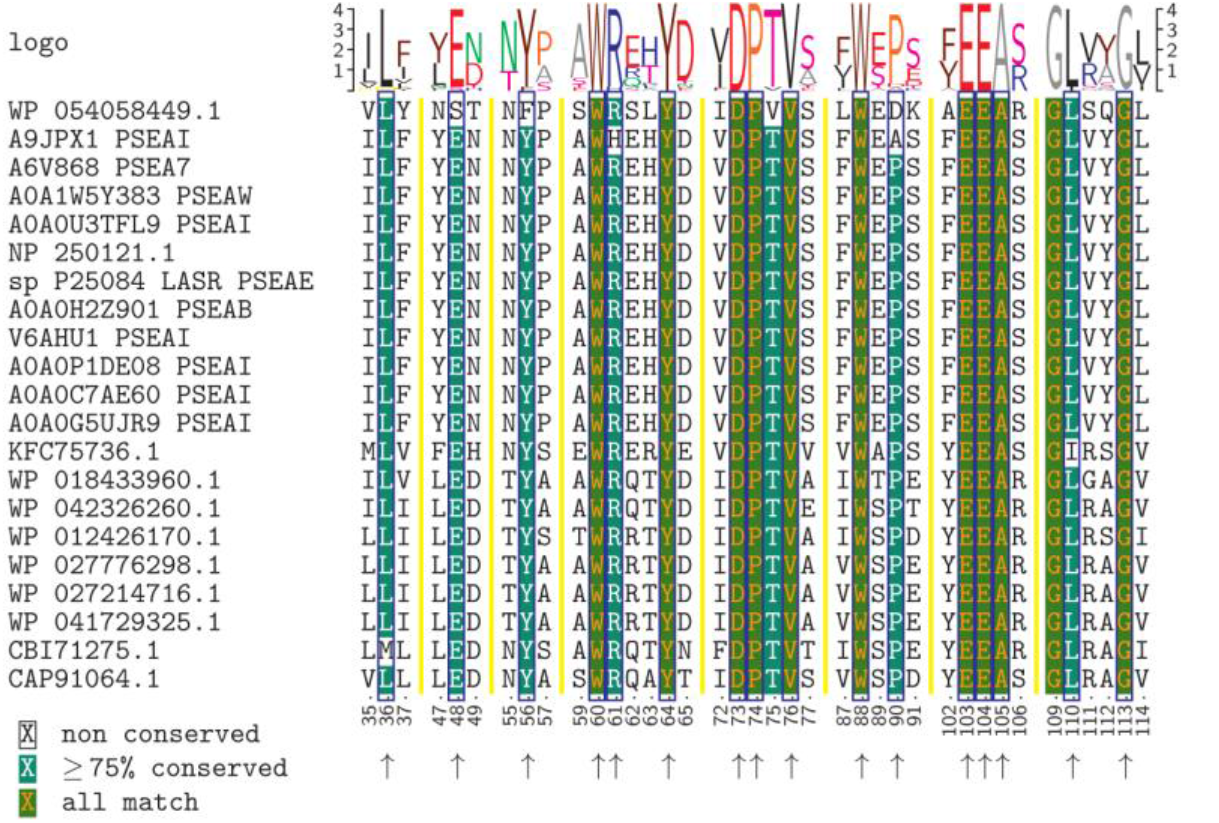
Interaction of conserved amino acids of LBD of LasR with the autoinducer molecule. Blue boxes and arrows point the amino acid residues that interact with 3-0-C12-HSL.

The new binding state with LBD-SLR-DBD bridge or “the bridge” of LasR with energy ~-166.456 kJ/mol (Table 2), where residues Leu236, Leul77, Vall76, Phe219, Lysl82 and Trpl9 contribute most (Figure 19).

**Fig. 17.**
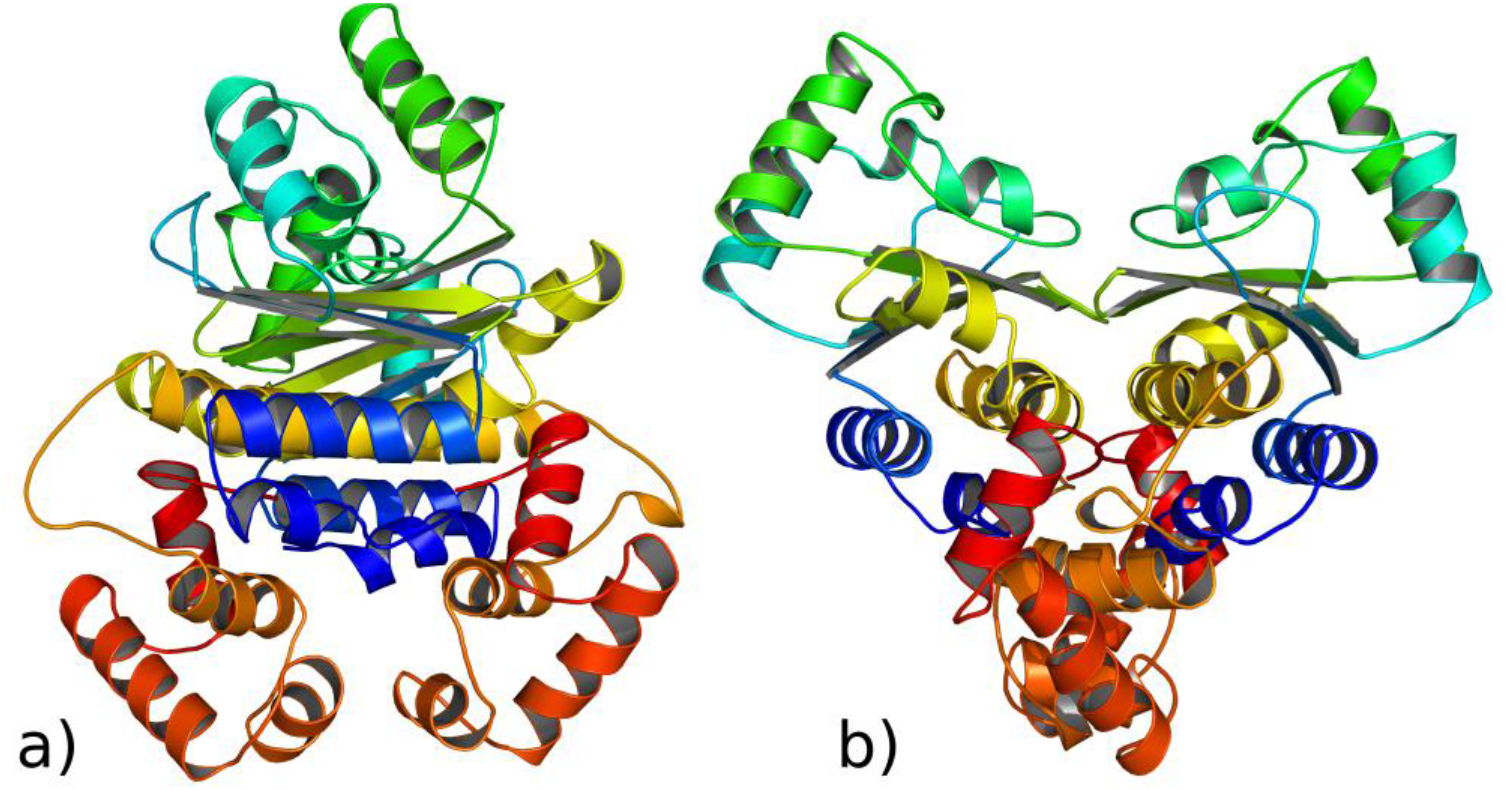
LasR monomer-LasR monomer docking result based on centroid conformation from cluster 2 of simulation 2 (3-0-C12-HSL with LBD) a) left view b) front view’.

For simulation 3 docking the top model contains 80 Members and the scores for the docking model were -951.4 for the Center and -1332.0 and for the lowest energy, thus suggesting a favourable binding mode (Figure 18).

**Fig. 18.**
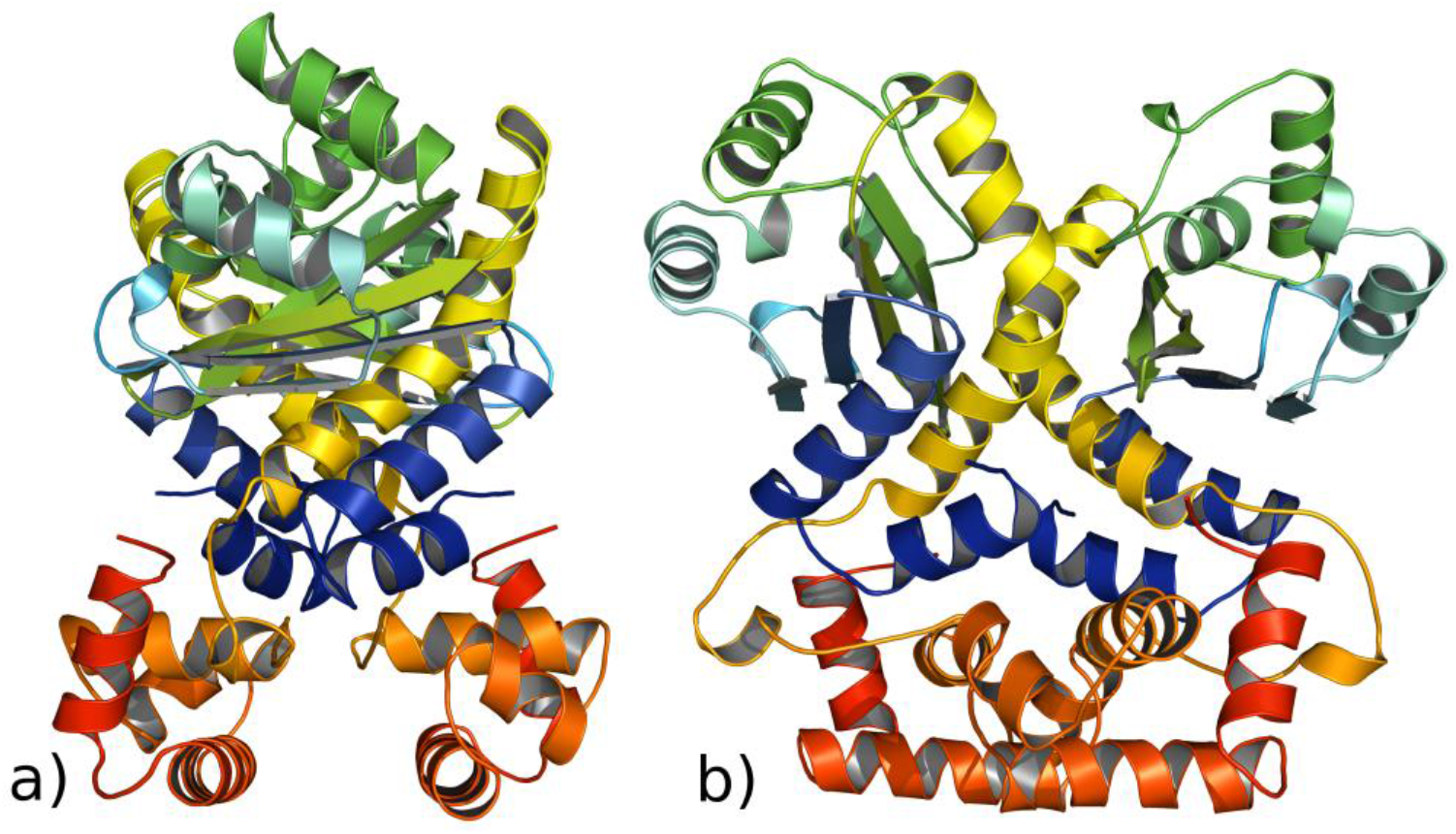
LasR monomer-LasR monomer docking result based on centroid conformation from cluster 1 of simulation 3 (3-0-C12-HSL with the “bridge”) a) left view b) front view.

**Fig. 19.**
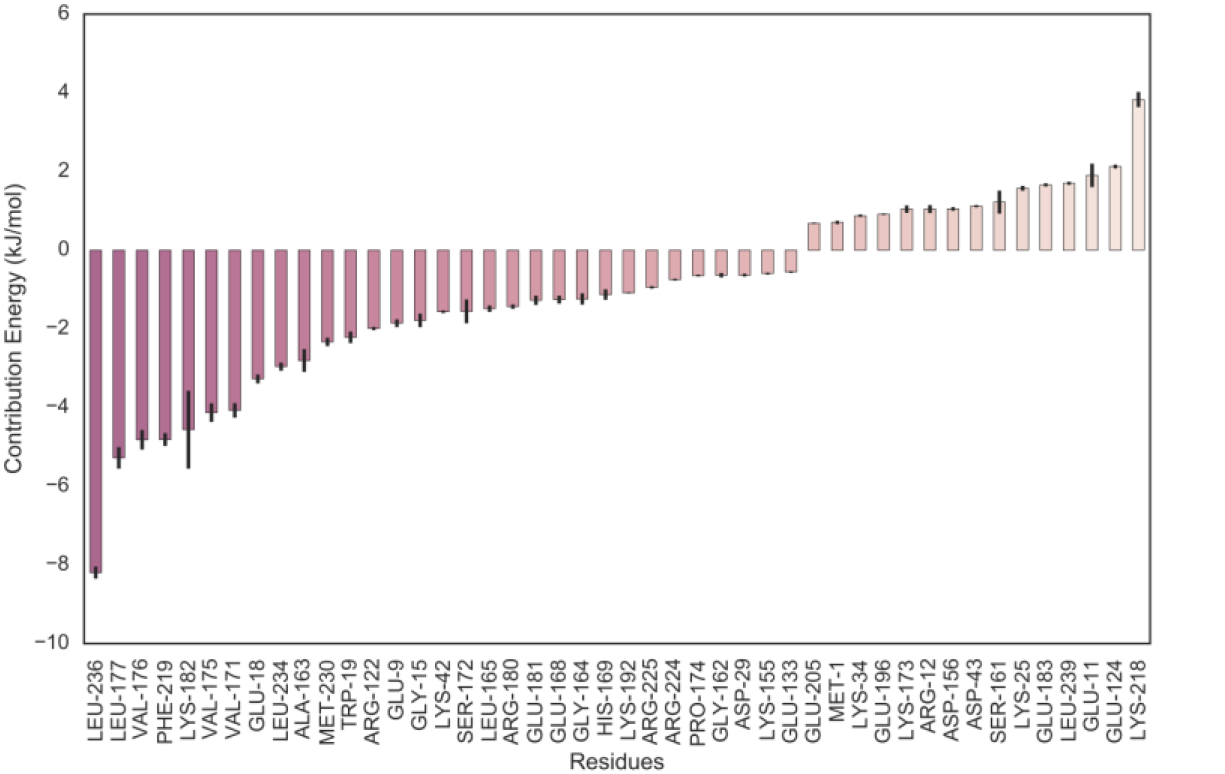
Energy contribution of LBD-SLR-DBD bridge amino acid residues from simulation 3.

Analysis of residue sequence alignment show that 3-0-C12-HSL interacts with very conservative amino acids such as Leu236, Leul77, Phe219, Trpl9 (Fig. 19, 20). 3-0-C12-HSL interacts with 12 fully conserved amino acids. In total 16 amino acids participate in the interaction, where amino acid conservation is more than 75%.

**Fig. 20.**
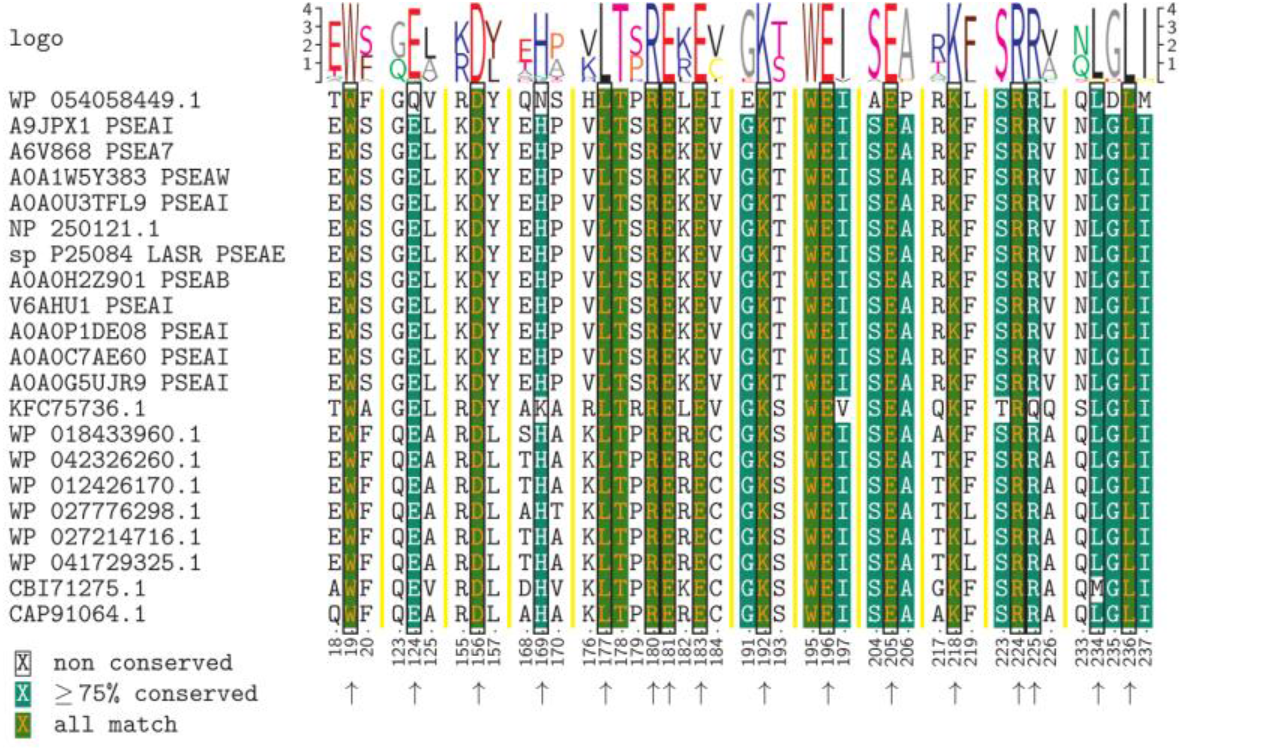
Interaction of conserved amino acids of LBD-SLR-DBD bridge (“the bridge”) with the autoinducer molecule. Black boxes and arrows point the residues that interact with 3-0-C12-HSL.

In this new binding site Trpl9. Aspl56 from LBD. Leul77 from SLR and Argl80. Glul 81. Glul96, Lys218, Arg224. Lys236 of DBD participate in the interactions with LasR. Lactone head group interacts with conserved Trpl9 of LBD. This result clearly suggests that both of the C-terminal and the N-terminal of LasR interact with 3-0-C12-HSL.

## Protein Docking of MD models

It is known that LasR binds to the corresponding promoter DNA region as a dimer [1] after interacting with the autoinducer. Thus, the monomeric LasR-3-0-C12-HSL complexes were subjected to dimerization. The protein docking experiments using structures from MD runs were performed with ClusPro 2.0 [67, 77-79] because of its success in the CAPRI (Critical Assessment of Predicted Interaction). Each centroid conformation was extracted from simulations 2 and 3 and was used for docking. For the selection of the model, we used the approach as recommended by the authors of ClusPro [67], suggesting to find the most populated clusters.

From simulation 2 the top model contains 122 members and the scores for the docking model were -1440.7 for the centre and -1517.9 for the lowest energy, suggesting a favourable binding mode (Figure 17).

The interaction of 3-0-C12-HSL with the “bridge” creates a possible conformation for dimerization. From our analysis, it is observed that 3-0-C12-HSL has two binding modes to LasR protein. The 3-0-C12-HSL, AI ligand (Figure 1) having a long hydrophobic tail is capable of binding both to LBD and to the “bridge” of LasR. The binding of 3-0-C12-HSL provokes conformational transitions of the transcriptional regulator LasR. The two binding conformations are capable of dimerization.

## Conclusion

From the simulations, it can be safely concluded that the AI ligand 3-0-C12-HSL, can bind both to LBD and to “bridge” of transcriptional regulator LasR. This suggests that there are multiple binding modes rather than one. The interaction with the LBD-SLR-DBD bridge is a novel site. The analysis of binding energy shows that the interaction of 3-0-C12-HSL with LDB-SLR-DBD bridge is stronger than with LBD. Conservative amino acids such as Leu236, Phe219, Leul77, Lysl82 and Trpl9, contribute most during the interaction with LBD-SLR-DBD bridge. This could suggest that for the DNA binding capability, it is necessary the interaction of 3-0-C12-HSL with the “bridge”.

Both conformational transitions favour dimerization, so this raises the question what would be the roles of different binding sites. This part needs further research. This study may reveal new insights of the interactions of the native 3-0-C12-HSL ligand with transcriptional regulator LasR of P. aeruginosa. Results from this study may be used for future drug development endeavours.

## Additional files

### Additional file 1

**Figure SI.** PROCHECK plot of LasR model. **Figure S2.** PROCHECK statistics of LasR model. **Figure S3.** Verify3D score plot of LasR model. **Figure S4.** Determination of exhaustiveness after PCA and cluster analysis. **Figure S5.** Exhaustiveness clusterization quality analysis. **Figure S6.** RMSD evolution through time for LasR monomer structure. The RMSD is calculated for the backbone atoms of the residues. **Figure S7.** Rg evolution through time for LasR monomer structure. The Rg is calculated for the backbone atoms of the residues. **Figure S8.** Agglomerative clustering with a different number of clusters for the LasR monomer run. In this case, a cluster count of **4** is the best option. **Figure S9.** The plot of chemical shifts of Cα-atom vs. residue number. **Figure S10.** Secondary structure analysis of LasR protein from simulation representative structure using PDBSum. **Figure Sll.** K-means clustering with a different number of clusters for the 3-0-C12-HSL docking data. In this case, a cluster count of 3 is the best option. **Figure S12.** Mean Autodock Vina binding energy score for cluster 1 and percentage from the total of conformations. **Figure S13.** Mean Autodock Vina binding energy score for cluster 2 and percentage from the total of conformations. **Figure S14.** Mean Autodock Vina binding energy score for cluster 3 and percentage from the total of conformations. **Figure S15.** Principal component analysis of ligand (3-0-C12-HSL) conformations obtained from various docking programs. **Figure S16.** PCA and RMSD analysis for simulation 2 of docking cluster 2. Left) PCA analysis Right) Colour-coded RMSD of the simulation obtained from Ward-linkage cluster analysis. **Figure S17.** The frequency of hydrogen bonds of cluster 2 in simulation 2 **Figure S18.** Hydrogen and hydrophobic interactions from centroid snapshot of cluster 2 in simulation 2 using LigPlot+. **Figure SI9.** PCA and RMSD analysis for simulation 3 of docking cluster 3. Left) PCA analysis Right) Colour-coded RMSD of the simulation obtained from Ward-linkage cluster analysis. **Figure S20.** The frequency of hydrogen bonds of cluster 1 in simulation 3. **Figure S21.** Hydrogen and hydrophobic interactions from centroid snapshot of cluster 1 in simulation 3 using LigPlot+. (DOCX)

**Additional file 2:** Movie of LasR monomer MD simulation. (MP4)

**Additional file 3:** Movie of MD simulation 1. (MP4)

**Additional file 4:** Movie of MD simulation 2. (MP4)

**Additional file 5:** Movie of MD simulation 3. (MP4)

**Additional file 6:** Full sequence alignment of LasR using ClustalW algorithm. (PDF)

**Additional file 7:** Full sequence alignment of LasR using Clustal Omega algorithm. (PDF)

**Additional file 8:** Full sequence alignment of LasR using Muscle algorithm. (PDF)

## Declarations

### Acknowledgments

We thank Yerevan Physics Institute for providing time on the cluster for the molecular dynamics simulations.

### Availability of data and materials

The analysed datasets are available as additional files.

### Authors’ contributions

HG designed the construct and simulations and performed the analyses. HG ST wrote the manuscript. HV ST LH supervised this work. All authors have reviewed and approved the final manuscript.

### Consent for publication

Not applicable.

### Ethics approval and consent to participate

Not applicable.

